# Parasite-Induced IFN-γ Regulates Host Defense via CD115 and mTOR-Dependent Mechanism of Tissue-Resident Macrophage Death

**DOI:** 10.1101/2023.06.23.546211

**Authors:** Andrew T Martin, Shilpi Giri, Alexandra Safronova, Sophia I Eliseeva, Samantha F Kwok, Felix Yarovinsky

## Abstract

Host resistance to a common protozoan parasite *Toxoplasma gondii* relies on a coordinated immune response involving multiple cell types, including macrophages. Embryonically seeded tissue-resident macrophages (TRMs) play a critical role in maintaining tissue homeostasis but their role in parasite clearance is poorly understood. In this study, we uncovered a crucial aspect of host defense against *T. gondii* mediated by TRMs. Through the use of neutralizing antibodies and conditional IFN-γ receptor-deficient mice, we demonstrated that IFN-γ directly mediated the elimination of TRMs. Mechanistically, IFN-γ stimulation rendered macrophages unresponsive to macrophage colony-stimulating factor (M-CSF) and inactivated mTOR signaling by causing the shedding of CD115 (CSFR1), the receptor for M-CSF. Further experiments revealed the essential role of macrophage IFN-γ responsiveness in host resistance to *T. gondii.* The elimination of peritoneal TRMs emerged as a host defense mechanism aimed at limiting the parasite’s reservoir. The identified mechanism, involving IFN-γ-induced CD115 and mTOR-dependent cell death, provides insights into the adaptation of macrophage subsets during infection and highlights a crucial aspect of host defense against intracellular pathogens.

## INTRODUCTION

Macrophages play a central role in detecting and eliminating bacterial, parasitic, and viral pathogens^1, 2^. They are uniquely positioned to sample the environment in order to detect pathogens, dead cells, and inflammatory mediators for the initiation of immune responses and tissue repair ^3–9^. There are two major types of macrophages, referred to as tissue-resident (TRM) and monocyte-derived macrophages^10–13^. TRMs, such as large peritoneal macrophages (LPMs) and Kupffer cells are embryonically derived and are maintained through self-proliferation independently from the monocyte-derived cells^14–17^. M-CSF signaling through activation of its receptor, CSFR1 (CD115), is required for both TRM development and maintenance in the peripheral tissues^18–21^. In contrast, monocyte-derived macrophages are frequently associated with both protective and pathological inflammatory responses seen during microbial infections^4, 22^. The requirement of M-CSF for the monocyte-derived macrophages is more complex; a complete deficiency in CD115-signaling results in their developmental defects while transient or partial M-CSF inactivation has a limited role on the frequency or the absolute numbers of monocyte-derived macrophages^8, 18–20^.

Monocyte-derived macrophages play a central role in pathogen elimination via multiple mechanisms including production of proinflammatory cytokines, nitric oxide, and reactive oxygen species^3, 4, 7^. Monocytes and monocyte-derived macrophages closely cooperate with dendritic cells in the regulation of T cell responses^23–25^. In contrast to monocyte-derived macrophages, our knowledge of TRM functions in the context of microbial infection is relatively limited. The unique response of TRMs to microbial infection is called the macrophage-disappearance reaction (MDR) which is characterized by contraction of the TRM population due to poorly understood mechanisms including tissue adherence, migration, and cell death during the first phase of the inflammatory response^26, 27^. Following their diminution during microbial or viral insult, the remaining TRMs undergo M-CSF-dependent proliferation, which contributes to their repopulation. This repopulation is bolstered by recruited monocytes that acquire a TRM-like phenotype during the resolution phase of the inflammatory response. Being located in a sterile environment, peritoneal macrophages serve as an excellent experimental model to study the first response of TRMs to microbial infections. Peritoneal macrophages are predominantly comprised of TRMs that can be identified as CD11b+F4/80+CD102+ cells expressing low levels of MHCII in naïve mice^28^. Monocyte-derived macrophages represent a minor subset and express low levels of F4/80 and CD102, and high levels of MHCII. Here we investigate the dynamic response of the peritoneal TRMs to *T. gondii*, a common intracellular pathogen that triggers a highly polarized Th1 response^29, 30^. We observed that *T. gondii* infection and the associated immune response results in elimination of TRMs in peritoneal cavity, liver, and intestine during the acute response to the parasitic infection. We established that the loss of TRMs is mediated by IFN-γ that is signaled directly to the macrophages and resulted in the shedding of CD115 from the macrophage cell surface. Our additional experiments revealed that the loss of CD115 triggered the cell death of TRMs via an mTOR-dependent mechanism that we recently identified in the context of the epithelial cell responses to IFN-γ^31^. Additional experiments uncovered that this TRM response to IFN-γ was essential for host resistance to the parasite, and the preservation of TRMs achieved by selectively targeting the IFN-γ receptor in TRMs caused an increased parasite burden responsible for the acute susceptibility of infected mice to *T. gondii* infection. When combined, our results established that IFN-γ was required and sufficient for triggering the CD115 and mTOR-dependent mechanism of LPM death, which played a role for host resistance to *T. gondii* by limiting a reservoir for the common intracellular parasite.

## RESULTS

### Loss of tissue-resident macrophages during *T. gondii* infection in vivo

Macrophages are sentinels of the immune system and are present in all lymphoid organs and tissues. These cells are involved in early interactions with pathogens, including *T. gondii* ^25, 32–34^. Two distinct macrophage subsets coexist in the peritoneal cavity^28^. The LPMs are seeded during embryonic development and represent the dominant population in naïve mice (Supplemental Fig. 1 and Fig. 1). LPMs are characterized by high levels of the canonical surface markers CD11b and F4/80 (Fig 1A and Supplemental Fig. 1). The LPMs can also be identified as Lin-CD11b+CD102+ cells that express low levels of MHCII (Fig. 1C and Supplemental Fig. 1). In contrast, monocyte-derived small peritoneal macrophages (SPMs) express low levels of F4/80 and CD102, and high levels of MHCII and represent a minor population in naïve mice (Fig 1A, 1B, and Supplemental Fig. 1). It has been established that monocytes and monocyte-derived macrophages play a crucial role in host defense against the intracellular parasite *T. gondii*^30, 32, 35^; but the roles of resident macrophages in controlling the immune response during infection with *T. gondii* is largely unknown.

**Figure 1.**
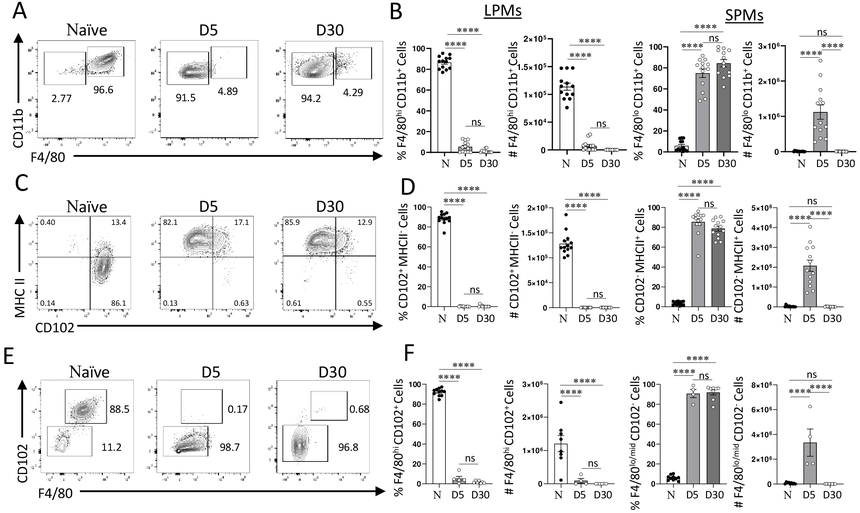
Resident peritoneal macrophages are lost during systemic *T. gondii* infection. (A) Flow cytometric analysis of large (CD11b+F4/80+) and small (CD11b+F4/80-) peritoneal macrophages measured on days 5 and 30 post intraperitoneal infection with 20 cysts of ME49 *T. gondii* per mouse. (B) Relative frequencies and absolute cell numbers of LPMs and SPMs measured on days 5 and 30 post intraperitoneal infection with *T. gondii*. (C-D) An alternative flow cytometric analysis of LPMs (CD11b+CD102+MHCII-) and SPMs (CD11b+CD102-MHCII+) in peritoneal cavity on days 5 and 30 post *T. gondii* infection. (E-F) Quantifications of LPMs (CD11b+CD102+F4/80+) and SPMs (CD11b+CD102-F4/80-) using the alternate gating strategy on days 5 and 30 post *T. gondii* infection. The results are representative of at least five independent experiments. Error bars = mean ± SEM ns P > 0.05, **** P < 0.00001.

We observed major changes in the composition of macrophage subsets during the acute response to *T. gondii.* We observed a nearly complete loss of the LPMs by day 5 post-infection (Fig. 1). This observation was established independently of the gating strategies used to define the LPMs as either CD11b+F4/80+ (Fig. 1A, 1B) or CD11b+CD102+MHCII-cells (Fig. 1C-1F), strongly suggesting that the acute response to *T. gondii* led to LPM loss (Fig. 1). In striking contrast, SPMs became the dominant macrophage subset during the acute response to the parasite. This was evident from the analysis of the absolute and relative frequencies of the SPMs on day 5 post infection (Fig. 1B, 1D, and 1F). While massive infiltration of monocyte-derived macrophages to the site of *T. gondii* infection has been previously described^34, 36–38^, the loss of the LPMs was unexpected and prompted us to further investigate the response of this macrophage subset to the parasite. We observed that even during the chronic stage of infection, LPMs were practically undetectable in the peritoneal cavity of the infected mice, indicating the parasitic infection caused a permanent change in the macrophage subsets (Fig. 1). Not only was there a dramatic reduction in the relative frequencies of LPMs, which could be explained by the migration of inflammatory monocytes, but an absolute quantification of LPMs revealed a complete or nearly complete loss of this macrophage subset that persisted past the acute stage of infection (Fig. 1B, 1D, and 1F).

Furthermore, experimental peritoneal infection resulted in a near complete elimination of hepatic resident macrophages, Kupffer cells (KCs), that were identified via VSIG4 (also known as CRIg) that is selectively expressed by KCs (Supplemental Fig. 2A). The disappearance of KCs coincided with the loss of LPMs (Supplemental Fig. 2B).

We also discovered that, similar to the experimental intraperitoneal *T. gondii* infection, the natural oral route of infection led to the loss of TRMs in the intestine (Fig. 2A-2C). Interestingly, oral infection with *T. gondii* also resulted in the loss of LPMs (Fig 2D, 2E), but it did not lead to significant recruitment of SPMs to the peritoneal cavity (Fig. 2E). These findings are significant as they indicate that the recruitment of SPMs and the loss of LPMs are distinct events in immune responses to *T. gondii*. While the mechanism of SPM recruitment to the site of infection has been previously established as a CCL2 dependent process^39–42^, the loss of TRMs regardless of the infection route, prompted us to investigate the underlying mechanism behind TRM depletion during acute responses to the parasite.

**Figure 2.**
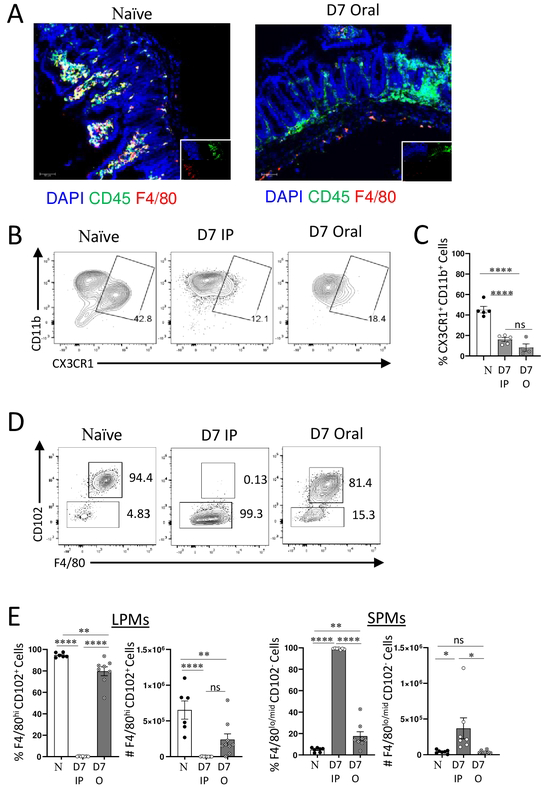
Resident intestinal and peritoneal macrophages are lost during oral *T. gondii* infection. (A) Representative immunohistochemical and (B) flow cytometric analysis of small intestinal macrophages defined as CD45+F4/80+CXCR1+ cells on day 7 post oral or intraperitoneal (IP) infection with 20 cysts of ME49 *T. gondii*. DAPI was used as nuclear stain. Original magnification 200X. (C) Frequency of CD45+CX3CR1+CD11b+ cells from the lamina propria on day 7 following intraperitoneal (D7 IP) or oral (D7 Oral) infection with 20 cysts of ME49 *T. gondii*. (D) Flow cytometric analysis of large (CD11b+F4/80+) and small (CD11b+F4/80-) peritoneal macrophages measured on day 7 post IP or oral infection with 20 cysts of ME49 *T. gondii.* (E) Quantifications of peritoneal LPMs (CD11b+CD102+F4/80+) and SPMs (CD11b+CD102-F4/80-) on day 7 post IP or oral infection with *T. gondii.* The results are representative of three independent experiments. Error bars = mean ± SEM, ns P > 0.05, * P < 0.01, ** P < 0.00, **** P < 0.00001

### IFN-γ is required and sufficient for the loss of large peritoneal macrophages

IFN-γ is a crucial cytokine that regulates both host resistance and immunopathological responses to *T. gondii* infection^43–46^. IFN-γ mediated activation of macrophages, known as priming, is required for the elimination of intracellular pathogens including *T. gondii*^7, 30, 44, 47, 48^. At the same time, the results from our and other laboratories revealed that IFN-γ can trigger major changes in tissue cellular composition via triggering cell death during an inflammatory response^31, 49–53^. To determine if IFN-γ played a role in the observed loss of LPMs seen in *T. gondiii* infected mice, we first analyzed macrophages in parasite-infected mice in the presence of IFN-γ blocking antibodies. We observed that neutralization of IFN-γ prevented the loss of LPMs during the acute response to the parasite (Fig. 3A, 3B). These results suggested that IFN-γ was required for the disapperance of LPMs seen in *T. gondii*-infected mice (Fig. 3A, 3B).

**Figure 3.**
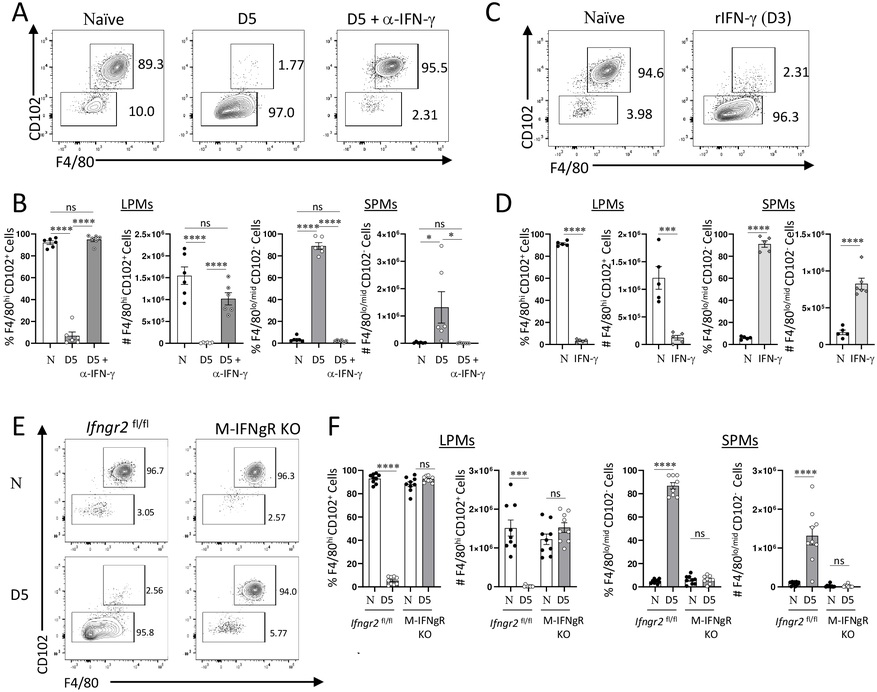
IFN-γ mediates loss of LPMs *in* vivo. (A) Flow cytometric analysis of LPM (CD11b+F4/80+CD102+) and SPM (CD11b+F4/80-CD102-) populations from naïve, *T. gondii* infected, and *T. gondii* infected mice treated with 200 μg anti-IFN-γ antibody on day 5 post intraperitoneal infection. (B) Quantification of LPM and SPM frequencies and absolute numbers in mice infected with *T. gondii* in the presence or absence of IFN-γ neutralizing antibody on day 5 post intraperitoneal infection with the parasite (20 cysts per mouse of the ME49 *T. gondii*). (C) Flow cytometric analysis of LPM and SPM populations in mice treated with recombinant IFN-γ (200 μg per mouse) for 72 h. (D) Quantification of LPM and SPM frequencies and absolute numbers in mice treated with recombinant IFN-γ (200 μg per mouse) for 72h. (E) Flow cytometric analysis of LPM (CD11b+F4/80+CD102+) and SPM (CD11b+F4/80-CD102-) populations in naïve and *T. gondii* infected MLys-Cre x *Ifngr2*^flox/flox^ (M-IFNgR KO) and *Ifngr2*^flox/flox^ mice on day 5 post intraperitoneal infection with 20 cysts per mouse of the ME49 *T. gondii*. (F) Quantification of LPM and SPM frequencies and absolute numbers in *T. gondii* infected MLys-Cre x *Ifngr2*^flox/flox^ (M-IFNgR KO) and *Ifngr2*^flox/flox^ mice on day 5 post intraperitoneal infection. The results are representative of three independent experiments. Error bars = mean ± SEM, ns P > 0.05, * P < 0.01, ** P < 0.00, *** P < 0.0001, **** P < 0.00001.

We next examined whether IFN-γ was sufficient for the elimination of LPMs *in vivo* in the absence of parasitic infection. This was crucial as *T. gondii* frequently infects macrophages and can modulate their functions via multiple virulence factors injected into the infected cells^54–58^. To test this possibility, naïve mice were injected with recombinant IFN-γ (rIFN-γ) and the macrophage subsets in the peritoneal cavity were analyzed 3 days later. We observed that injection of rIFN-γ was sufficient to drive the loss of LPMs (Fig. 3C, 3D). These results strongly indicated that even in the absence of *T. gondii* infection, IFN-γ can trigger the dissaperance of LPMs. Of note, the experimental dose of rIFN-γ mirrored the levels of circulating IFN-γ seen in the infected mice, emphasizing the physiological relevance for this cytokine treatment (not shown).

When combined, our results formally established that IFN-γ was both required and sufficient for the changes in peritoneal macrophage composition characterized by the loss of LPMs (Fig. 3A-3D).

### IFN-γ signaling in macrophages is required for the loss of large peritoneal macrophages

IFN-γ has pleiotropic effects on multiple cell types^59^. Experiments with systemic blockage of or supplementation with rIFN-γ do not discriminate between the direct effects of this cytokine on peritoneal macrophages or indirect effects caused by IFN-γ activating other cell types (Fig. 3A-3D). This knowledge is crucial for deciphering the molecular mechanisms required for the loss of LPMs during *T. gondii* infection.

To examine whether the direct effects of IFN-γ on macrophages are responsible for LPM loss during the acute response to *T. gondii*, mice with an IFN-γ receptor deficiency specific to myeloid cells were generated by crossing Mlys-cre mice with *Infgr2*^flox/flox^ mice. Myeloid-cell-specific IFN-γRII knockout mice (M-IFNgR KO) and their cre-negative littermate controls were next infected with the parasite and changes in macrophage populations were analyzed on day 5 post-infection (Fig. 3E, 3F). As anticipated, cre-negative mice were indistinguishable from the wild-type (WT) controls and lost their LPMs during the acute response to *T. gondii* (Fig. 3E, 3F). In contrast, we observed that both the frequencies and absolute numbers of LPM and SPM populations were practically unchanged when comparing day 5 infected and naïve control M-IFNgR KO mice (Fig. 3E, 3F). These results indicate that IFN-γ acting directly on peritoneal macrophages is responsible for the loss of LPMs (Fig. 3E, 3F).

### IFN-γ treatment drives the death of large peritoneal macrophages

Several non-mutually exclusive mechanisms can explain the rapid loss of LPMs during the acute response to *T. gondii*. Among the major possibilities are migration of LPMs caused by *T. gondii* infection, or IFN-γ mediated death of LPMs, a mechanism recently reported for intestinal epithelial cells^31, 49^.

We first examined whether IFN-γ can trigger LPM migration to the omentum. The omentum is located in the peritoneal cavity and iscomposed of two mesothelial sheets enclosing adipose tissue. Recenly, it was observed that *T. gondii* infected macrophages and dendritic cells can rapidly relocate from the peritoneum to the omentum for induction of T cell responses to the parasite^60^. To minimize artifacts associated with cell isolation or incomplete analysis of the omentum, the entire omenta from naïve and *T. gondii* infected mice were next analyzed by whole-mount imaging (Supplemental Fig. 3). While an increase in CD11b+ cells was observed in the omentum analyzed from *T. gondii* infected mice, only an incremental amplification in peritoneal LPM-specific CD102+ cells^61, 62^ was seen (Supplemental Fig. 3). These results indicated that the loss of LPMs during parasitic infection was not the result of cell relocation from the peritoneal cavity to the omentum.

To test a hypothesis that *T. gondii*-induced IFN-γ lead to the death of LPMs via a mechanism recently reported by us for Paneth cells^31^, we first applied an *in vitro* system with peritoneal LPMs stimulated with rIFN-γ (Fig. 4A, 4B). Timelapse imaging analysis of macrophages treated with rIFN-γ was performed for 18 hours and dying cells were detected with propidium iodide (PI) staining (Fig.4A). We observed that IFN-γ treatment resulted in the rapid appearance of PI+ cells (red) during the course of the experiment (Fig. 4A). In contrast, only minimal cell death was seen in the untreated controls. An independent quantitative imaging experiment confirmed the initial observation and revealed that IFN-γ triggered the death of LPMs (Fig. 4B). These results revealed that IFN-γ can trigger the death of LPMs, but does not formally demonstrate the direct causality of this cytokine in the induction of macrophage death. To examine the direct effect of IFN-γ signaling on macrophages, we isolated macrophages from the peritoneal cavity of M-IFNgR KO mice and treated them with rIFN-γprior to utilizing imaging flow cytometry to assess cellular death. We observed that a lack of responsiveness to IFN-γ precluded the death of LPMs (Fig. 4C, 4D). At the same time and analogous to the WT macrophages, the majority of the cre-negative *Ifngr*^flox/flox^ macrophages were PI+ within 18 hours of treatment with rIFN-γ (Fig. 4C, 4D). When combined, these results revealed that IFN-γ signaling in macrophages directly leads to the loss of LPMs due to their death.

**Figure 4.**
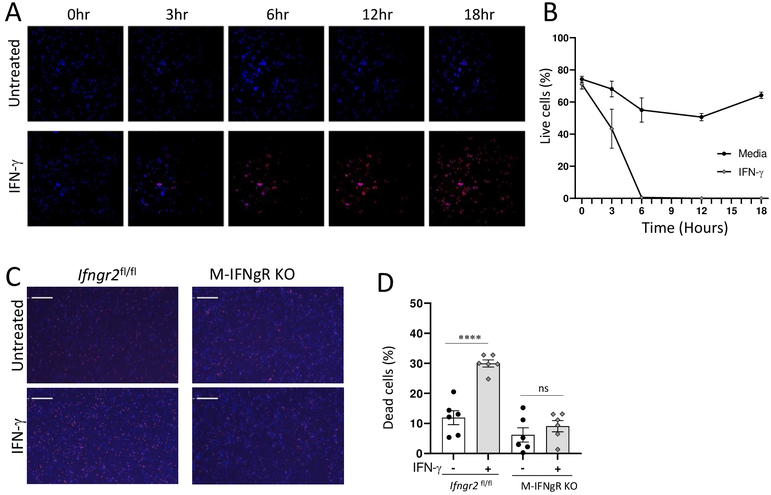
IFN-γ mediates the death of LPMs. (A) LPMs were cultivated in a temperature, CO_2_, and humidity-controlled chamber in the presence or absence of rIFN-γ. Time lapse imaging was performed for 18 h in the presence of Hoescht 3364 and Propridium Iodide (PI), and the representative snapshots at 3, 6, 12, and 18h are shown. (B) Quantification of live LPMs detected as PI-negative cells at 3, 6, 12, and 18 h post treatment with rIFN-γ. (C) LPMs prepared from M-IFNgR KO mice or the appropriate WT controls (Cre-negative *Ifngr2*^flox/flox^ littermates) were plated onto black walled, flat-bottom, 96 well plates and the cells were either left untreated or treated with rIFN-γ for 18 hours. Images were captured using the Celigo imaging cytomer and (D) dead cells were quantified as PI+ cells after 18 h. Scale bar = represents 200 μm. Error bars = mean ± SEM. ns P > 0.05, ****P < 0.00001.

### IFN-γ triggers a CD115- and mTOR-dependent death of large peritoneal macrophages

IFN-γ triggers profound metabolic changes in multiple cell types, including macrophages^63, 64^. One of the key molecules that senses cell metabolism is the mTORC1 kinase complex, which plays a critical role in integrating growth receptor signaling with cellular functions, including cell survival^65^. In addition, we have recently observed that in the context of *T. gondii* infection, IFN-γ leads to the death of intestinal Paneth cells via an mTOR-dependent mechanism distinct from apoptosis, necroptosis, or pyroptosis^31^. Furthermore, it has been demonstrated that IFN-γ can suppress the activity of mTORC1 in other cell types including macrophages^66^. Considering that, we next examined if mTOR supression was also responsible for the death of LPMs.

To test this hypotheiss, we first examined the requirements for mTORC1 activity in LPMs and SPMs. We observed that treatment of mice with the mTORC1 inhibitor rapamycin resulted in the rapid and selective loss of LPMs (Fig. 5A, 5B). While this treatment blocked mTOR activities in both LPMs and SPMs, only LPMs were susceptible to this treatment as evidenced by the analysis of dying cells in response to rapamycin treatment (Fig. 5C, 5D). We observed that only a small fraction of SPMs were stained with the dead cell reporter dye zombie yellow (Fig. 5C). Contrariwise, not only were practically all LPMs eliminated in rapamycin-treated mice, but the remaing LPMs were primarily positive for the dead cell dye (Fig. 5C, 5D.) These results implicated mTOR activity as a critical component in LPM survival.

**Figure 5.**
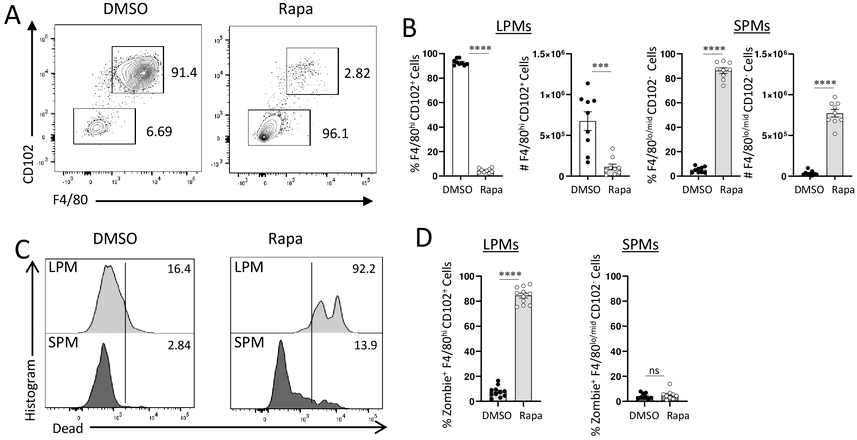
mTOR Inhibition leads to LPM death. (A) Flow cytometric analysis of large (CD11b+F4/80+CD102+) and small (CD11b+F4/80-CD102-) peritoneal macrophages isolated from WT mice treated with 300 μg rapamycin (Rapa) for 24 hours. (B) Relative frequencies and absolute cell numbers of LPMs and SPMs measured 24 h post injection of rapamycin. (C) Detection of dead or dying LPMs (Zombie Yellow+CD11b+F4/80+) and SPMs (Zombie Yellow+CD11b+F4/80-) 24 h post rapamycin treatment. (D) Relative frequencies of dead LPMs (Zombie Yellow+CD11b+F4/80+) and SPMs (Zombie Yellow+CD11b+F4/80-) 24 h post rapamycin treatment. The results are representative of three independent experiments. Error bars = mean ± SEM, ns P > 0.05, **** P < 0.00001

Among the various factors that are responsible for sustained mTORC1 activity in macrophages, the M-CSF-dependent activation of its receptor, CD115, is of particular interest. M-CSF signaling plays an indispensable role in the development of macrophages and is essential for the survival and maintenance of LPMs in the peritoneal cavity^9, 20^. For the above reasons, we examined the expression of CD115 in naïve or *T. gondii* infected mice, along with the animals stimulated with rIFN-γ. We observed that both parasitic infection and cytokine treatment *in vivo* resulted in the rapid loss of CD115 cell surface expression (Fig. 6A). Furthermore, when examining the peritoneal exudate fluid collected from the same mice, we observed the release of the extracellular portion of CD115 (Fig. 6B), suggesting that IFN-γ triggers shedding of CD115 from peritoneal macrophages.

**Figure 6.**
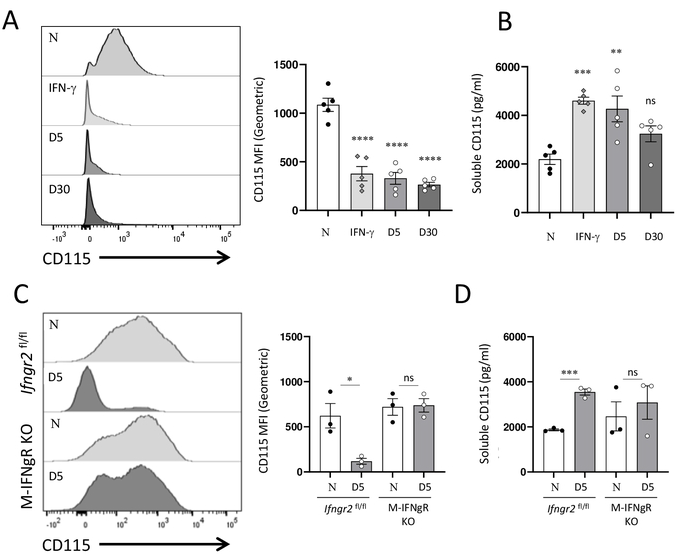
IFN-γ and *T. gondii* infection result in shedding of M-CSFR. (A) Flow cytometric analysis of CD115 expression on cell surface of peritoneal CD11b+ cells isolated from WT (C57BL/6) mice injected with rIFN-γ (200 μg per mouse, 72 h post treatment), or infected intraperitoneal with 20 cysts per mouse of the ME49 *T. gondii* for 5 and 30 days. The right panel shows quantification of CD115 MFI among CD11b^+^ cells. (B) Quantification of soluble CD115 detected in the peritoneal cavity of mice injected with rIFN-γ or infected with *T. gondii.* (C) Flow cytometric analysis of CD115 expression on cell surface of peritoneal CD11b+ cells isolated from M-IFNgR KO mice or their *Ifngr2*^flox/flox^ littermates infected intraperitoneal with *T. gondii* for 5 days. The right panel shows quantification of CD115 MFI among CD11b^+^ cells isolated on day 5 post infection with *T. gondii*. (D) Quantification of soluble CD115 detected in the peritoneal cavity of M-IFNgR KO mice or their *Ifngr2*^flox/flox^ littermate controls on day 5 post infection with *T. gondii.*. Error bars = mean ± SEM, ns P > 0.05, * P < 0.01, ** P < 0.00, *** P < 0.0001, **** P < 0.00001.

To determine if direct IFN-γ stimulation of macrophages leads to the loss of CD115 from the cell surface during *T. gondii* infection, CD115 expression was next analyzed in M-IFNgR KO mice infected with the parasite. We observed that the lack of IFNgR prevented the loss of CD115 cell surface expression in myeloid cells (Fig. 6C). In addition, the lack of IFNgR prevented the release of soluble CD115 into the peritoneal cavity of the infected mice (Fig. 6D). As anticipated, the appropriate cre-negative *IfngrII*^flox/flox^ mice behaved indistinguishable from the WT controls, and lost CD115 from the cell surface of the peritoneal macrophages (Fig. 6). When combined, these data revealed that IFN-γ leads to the shedding of CD115, an essential receptor for the survival and maintenance of LPMs, but not SPMs (Fig 6).

### Macrophage IFN-γ signaling is required for host survival during acute toxoplasmosis

The IFN-γ-mediated selective loss of LPMs in response to *T. gondii* infection prompted us to investigate whether this phenomenon is a host defense mechanism or an immunopathological response induced by IFN-γ. To distingushing among those possibilities, we first examined the survival of M-IFNgR KO mice infected with *T. gondii*. We observed acute susceptibility in these mice, displaying kinetics similar to those of complete IFN-γ-deficient mice (Fig. 7). These findings established that the macrophage response to IFN-γ is crucial for resistance against parasitic infection, even when other cell types can respond to IFN-γ (Fig. 7A). The acute susceptibility observed in the mice was directly associated with an uncontrolled parasite burden (Fig. 7B, 7C).

**Figure 7.**
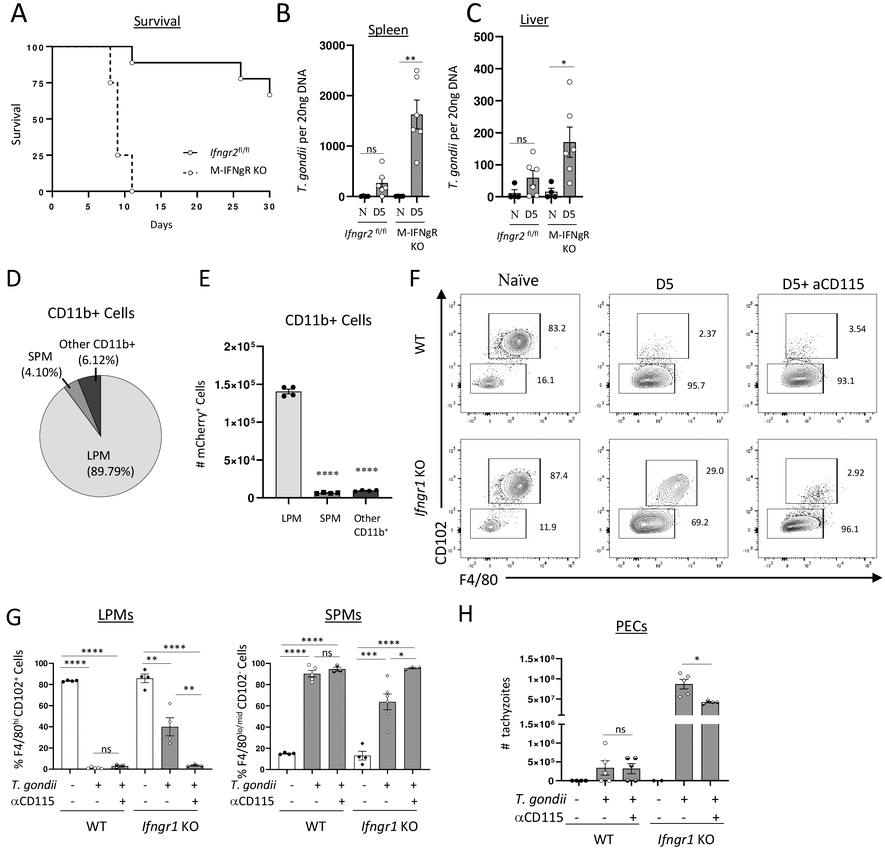
IFN-γ signaling in macrophages is required for host survival during toxoplasmosis. (A) Survival of M-IFNgR KO mice and their WT controls (Cre-negative *Ifngr2*^flox/flox^ littermates during intraperitoneal infection with 20 cysts of ME49 *T. gondii* from a combination of three experiments, each involving at least five mice per group. (B, C) Analysis of *T. gondii* parasite loads by qRT-PCR in (B) spleen and (C) liver. (D, E) Relative frequencies (D) and (E) absolute cell numbers of CD11b+ peritoneal cells infected in vitro with mCherry-expressing *T. gondii* (Pru strain) at 3:1 MOI for 3 h. (F) WT and *Ifngr1*^-/-^ mice were pretreated with 250ug of anti-CD115 or isotype control antibodies and intraperitoneally infected with 20 cysts of ME49 strain of *T. gondii* for 5 days. Representative flow cytometry plots and (G) quantification of LPM and SPM frequencies and absolute numbers in mice with the indicated treatments on day 5 post intraperitoneal infection. (H) Quantification of visible tachyzoites in the peritoneal cavity was performed on day 5 post infection. The results are representative of three independent experiments. Error bars = mean ± SEM, ns P > 0.05, * P < 0.01, ** P < 0.00, **** P < 0.00001

We next investigated whether the IFN-γ-dependent death of f LPMs limits the replication niche for the parasite. *In vitro* experiments revealed that *T. gondii* infection preferentially targets LPMs rather than SPMs (Fig. 7D, 7E), supportring a model that LPMs are an imporrtnat reservour for the parasite. To further investigate this in an in vivo setting, we employed a strategy to deplete LPMs by blocking the CD115 receptor (Fig. 7F). This approach allowed us to achieve the depletion of LPMs in an IFN-γ-independent manner. We observed that the depletion of LPMs through CD115 blockade had no impact on the parasite burden in wild-type (WT) mice, as they were already experiencing LPM loss due to IFN-γ, irrespective of the additional treatment (Fig. 7F, 7G, and 7H). However, when the response to IFN-γ was abolished, the depletion of LPMs resulted in a significant reduction in successful parasite replication (Fig. 7H). Although CD115-mediated depletion did not decrease the parasite burden in IFN-γ receptor-deficient mice to the level observed in WT controls (Fig. 7H), our experimental findings strongly support the hypothesis that the IFN-γ-mediated elimination of LPMs plays a pivotal role in host defense by controlling the elimination of a reservoir for the intracellular parasite *T. gondii* (Fig. 7H).

## DISCUSSION

Our study unveils a novel host defense mechanism against *T. gondii* infection, which involves the IFN-γ-mediated elimination of TRMs. We have demonstrated that IFN-γ directly impacts LPMs, leading to their death through CD115 and mTOR-dependent pathways. This host defense process plays a critical role in restricting the available niche for the intracellular parasite. Our findings expand upon the existing understanding of IFN-γ-mediated host resistance to intracellular pathogens, characterized by the activation of macrophages prior or during infection with *T. gondii*.

TRMs are recognized as key regulators in tissue homeostasis, organ development, and the resolution of infections. Although extensive research has been conducted on the development and lineage of TRMs using lineage tracing and RNAseq techniques^61^, their specific role in host defense remains incompletely understood. The loss of TRMs during microbial infection is a distinctive feature known as the MDR^27^. This response has been observed in relation to diverse microbial components, such as zymosan, LPS, and live bacterial pathogens, and the current study extends this concept to IFN-γ. Notably, the MDR or MDR-like loss of TRMs is not confined to the peritoneal cavity but has also been observed in other tissues. For instance, in the intestine, microbiota drives replacement of TRMs by monocyte-derived macrophages, suggesting that even low concentrations of microbial ligands can trigger the MDR^27^. Similarly, inflammatory responses can lead to the replacement of liver TRMs with monocyte-derived cells^67^, indicating that the loss of TRMs is a common physiological or pathophysiological response to microbial ligands and inflammatory cytokines.

In our investigation, we discovered that IFN-γ stimulation renders macrophages unresponsive to M-CSF by inducing the shedding of its receptor, CD115. Given the critical role of M-CSF in TRMs development and survival, the loss of CD115 is a crucial step contributing to the elimination of TRMs. M-CSF is a tighly controlled growth factor that contributes to monocyte to macrophage differentiation. Both macrophages and fibroblasts can prodcue M-CSF, and its levels regulate macrophage presence through autocrine and paracrine mechanisms^9, 68^. Thus, the regulation of M-CSF responsiveness is an effective strategy to control macrophage populations. Functionally, M-CSF can enhance phagocytic activity^69^, antigen presentation^70^, and cytokine production^71–73^, thereby augmenting the ability of macrophages to eliminate pathogens and modulate immune responses. The distinct requirement for mTOR signaling may suggest that monocyte-derived macrophages are better equipped to combat intracellular pathogens in the highly inflammatory environment caused by large amounts of IFN-γ seen during *T. gondii* infection. This may explain why TRMs cannot be reprogrammed by IFN-γ to acquire the functions of proinflammatory SPMs, although further investigation is needed to fully explore this hypothesis. Our study suggests that the IFN-γ-induced CD115 shedding is a rapid and efficient strategy to eliminate the preexisting TRMs. This mechanim of host defense cooperates with TLR-dependent and independent production of CCL2 that leads to the recruitment of monocytes to the site of infection that are indispensible for host resistance to *T. gondii*^39–42^. While the future biochemical studies are needed to define IFN-γ inducible enzymes responsible for CD115 shedding, it has been reported that metalloproteases, particularly ADAM17 (a disintegrin and metalloprotease 17) and ADAM10 (a disintegrin and metalloprotease 10) cleave the extracellular domain of CD115, releasing it as a soluble form into the extracellular environment^74–76^. These enzymes can be activated in response to various stimuli, including cytokines and microbial components^77^, suggesting a general strategy to regulate macrophage responses in different physiological and pathological conditions via regulation of responsivness to M-CSF.

In summary, our study reveals an IFN-γ-induced mechanism of host resistance to *T. gondii,* characterized by CD115 and mTOR-dependent macrophage death at the site of infection and potential sites of the parasite dissemination. Although additional studies are necessary to fully understand the necessity of TRM elimination, we speculate that it is a common host defense mechanism against intracellular infections. We provide experimental evidence supporting the concept that the elimination of TRMs and their subsequent replacement by monocyte-derived cells represents an IFN-γ-mediated host defense strategy against a common protozoan parasite capable of infecting macrophages.

## MATERIALS AND METHODS

### Mice

C57BL/6 mice were originally purchased from the Jackson Laboratories and maintained in the pathogen-free animal facility at the University of Rochester of Medicine and Dentistry. *Ifngr2^flox/flox^*mice were generated using targeted embryonic stem cells obtained from the Knockout Mouse Project repository and injected into C57Bl/6 albino blastocysts by the Fox Chase Cancer Center Transgenic Mouse Facility as previously described^31, 78^. Mlys-Cre (LysM-Cre, Strain #004781) mice were purchased from the Jackson laboratory. Mice for all experiments were age- and sex-matched. All animal experimentation in this study was reviewed and approved by the University of Rochester’s University Committee on Animal Resources (UCAR), the Institutional Animal Care and Use Committee (IACUC).

### Murine *T. gondii* infection and treatments

ME49 strain *T. gondii* tissue cyst (bradyzoite) stages were maintained through serial passage in Swiss Webster mice. For infections, brains of chronically infected Swiss Webster mice were mechanically homogenized by passage through a series of 18-gauge, 20-gauge, and 22-gauge needles. Experimental mice were intraperitoneally or orally infected with 20 *T. gondii* brain cysts (ME49 strain). In some experiments, mice were injected i.p. with 200 μg anti-IFN-γ (clone XMG1.2, BioXCell) on days 0, and 3 post *T. gondii* infection, with 200 μg of recombinant mouse IFN-γ alone (Genscript), 250 μg anti-CD115 (clone ASF98, BioXCell), or 300 μg Rapamycin (Sigma).

### Macrophage isolation and flow cytometric analysis

Peritoneal exudate cells (PECs) were isolated via lavage and single cell suspensions were briefly washed with PBS prior to red blood cell lysis using ACK lysis buffer. Prepared cells were stained with Zombie Yellow (BioLegend) to assess live status of cells. The following antibodies were used, though not necessarily all in the same panel: F4/80-BV785 (BioLegend; Clone BM8), CD115-BV421 (BioLegend; Clone AFS98), CD11c-BV421 (BD Bioscience; Clone HL3), CD115-BUV395 (BD Bioscience; Clone AFS98), CD19-BUV737 (BD Bioscience; Clone 1D3), CD226-PE (BD Bioscience; Clone TX42.1), CD11b-PE (eBioscience; Clone M1/70), MHCII (I-A/I-E)-PE/Cy7 (BioLegend; Clone M5/114.15.2), Ly6G-PE/CF594 (BD Bioscience; Clone 1A8), Ly6C-APC/eFluor780 (eBiosceince; Clone HK1.4), CD102-Alexa Fluor 647 (BioLegend; Clone 3C4 (mlC2/4)), CD102-Alexa Fluor 488 (eBioscience; Clone 3C4 [mlC2/4]), CCR2-FITC (BioLegend; Clone SA203G11), and CD11b-PerCP/Cy5.5 (BioLegend; Clone M1/70). Antibody cocktails were prepared in FACs staining buffer (phosphate-buffered saline, 1% FBS, 1% 0.5mM EDTA). Cellular fluorescence was measured using an LSRII Fortessa flow cytometer, and data were analyzed using FlowJo software (Tree Star).

### Visualization of macrophage death *in vitro*

For timelapse imaging, 2×10^5^ peritoneal macrophages per dish were imaged using a TE2000-U microscope (Nikon) coupled to a CoolSNAP HQ CCD camera with a 20x objective. Images were taken at the following intervals: brightfield images every 30 seconds and DAPI/Cy5 channels every 1 minute. Imaging took place over 18 hours in a humidified, temperature-controlled chamber maintained at 37°C for the duration of imaging. To assemble the timelapse move, images were compressed at normal quality and set at 100ms speed.

For quantitative analysis of cell death, 2×10^4^ freshly isolated PECs were plated in each well of a black-walled, clear bottom 96 well plates and macrophages were enriched for a final density of 1×10^4^ macrophages per well. Prior to imaging, an additional 100μl complete media was added to the wells containing either rIFN-γ or PBS (untreated) and a mixture of Hoescht 3364 and PI. Images were then captured using the Total (Brightfield) + Live (Blue-Hoescht) + Dead (Red-PI) setting on a Celigo Imaging Cytometer (Nexcelom).

### Determination of pathogen loads during infection

To determine *T*. *gondii* pathogen loads, total genomic DNA from animal tissue was isolated by using the DNeasy Blood and Tissue Kit (Qiagen) according to manufacturer’s instructions. PCR were performed by using SSOFast Eva Green Supermix (BioRad). Samples were measured by qPCR using a MyiQ Real-Time PCR Detection System (BioRad), and data from genomic DNA was compared with a defined copy number standard of the *T*. *gondii* gene *B1*.

### Immunofluorescence staining of the liver

For Immunofluorescence images, livers were fixed in 4% PFA in PBS at 4°C for 1 hour. Following fixation, samples were incubated in a 30% sucrose solution in PBS overnight at 4°C. Samples were then frozen in OCT (Tissue Tek) compound. 10 µm liver sections were washed three times with 50mM NH_4_Cl in PBS for 3 minutes prior to permeabilization with 0.25% Triton X-10 solution in PBS for 10 minutes at room temperature. Primary antibody staining for VSIG4 was performed in 2% serum in PBS overnight at 4°C. Slides were mounted in ProLong Gold (Molecular Probes). Specimens were imaged with a Leica SPE system (Leica DMi8) fitted with a Leica 63× objective NA 1.4.

### Identification of TRMs within the small intestine

For flow cytometric analysis of intestinal TRMs in the lamina propria, small intestine segments were washed, flattened, cut into 1.5cm pieces and shaken in 2 changes of HBSS with 5% FBS and 2 mM EDTA for 20 min at 37C . The suspension was passed through a mesh strainer to remove epithelial cells. The intestinal pieces were chopped using scissors and shaken in digestion media containing HBSS (Gibco) with 5% FBS, 1mg/ml Collagenase D, 2U/ml DNase I, 0.1U/ml Dispase for 30 mins at 37C. The digested samples were then vortexed briefly and passed through a 100 micron filter followed by staining for flow cytometry.

Cells were stained with the following fluorochrome-conjugated surface antibodies: CD45-BUV395 (BD Biosciences; Clone 30-F11), CD64-PE/Cy7 (Biolegend; Clone X54-5/7.1), CD11c-Alexa Fluor 488 (eBioscience; Clone N418), CD11b-APC/Cy7 (Biolegend; Clone M1/70), CX3CR1-BV785 (Biolegend; Clone SA011F11), and F4/80-APC (eBioscience; Clone BM8). Cellular fluorescence was measured using an LSRII Fortessa flow cytometer, and data were analyzed using FlowJo software (Tree Star).

### Whole mount immunofluorescence of the omentum

Mice were sacrificed and whole omenta were excised. Samples were placed in 6ml polypropylene tubes and blocked using FC Block (PharMingen, San Diego, CA) at 10μg/ml in 200μl of FACs buffer and placed on a shaker for 15 minutes at 4°C. Following incubation, samples were stained with the following antibodies for 2 hours at 4°C: CD11b-PE (eBioscience; Clone M1/70), CD102-Alexa Fluor 647 (BioLegend; Clone 3C4), and CD115-BV421 (BioLegend; Clone AFS98) or left unstained. Samples were washed twice by the addition of 4ml FACs buffer and rotated at 4°C for 30 minutes. After the final wash, samples were collected from the tubes using a snipped off P1000 tip and placed onto glass slides with FACs buffer. A cover slip was placed on top of the omentum and pressed down. Samples were viewed via fluorescence microscopy and digital images were acquired.

### Statistical analysis

All data were analyzed with Prism (Version 9.4.1; GraphPad, La Jolla, CA). These data were considered statistically significant when *P*-values were <0.05.

## Acknowledgments

This work was supported by NIAID Grants R01AI136538 and R01AI121090, and by the Burroughs Wellcome Foundation to F.Y. ATM was in part supported by the T32 training grant in Immunology, AI007285.

The authors are grateful to Dr. Minsoo Kim (University of Rochester) for help with the imaging, Dr. Taylor Ucello and Dr. Scott Gerber (University fo Rochester) for help with omentum isolation and imaging and to Dr. Yeojin Lee for KC analysis.

---

**Supplemental Figure 1.**
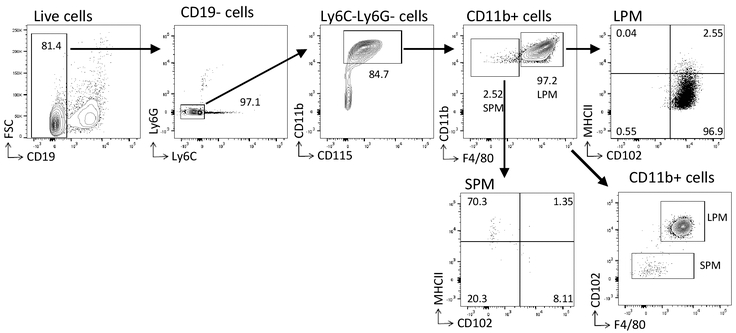
Peritoneal macrophage gating strategy. To identify peritoneal macrophage subpopulations, live (Zombie Yellow negative) single cells were gated for CD19-cells. LPMs were identified as F4/80+CD11b+ cells and SPMs were identified as F4/80-CD11b+ cells. LPMs and SPMs were additionally gated on CD102 and MHCII expression to validate the use of the alternate gating strategy for identifying LPMs as CD102+MHCII-cells and SPMs as CD102-MHCII+ cells.

**Supplemental Figure 2.**
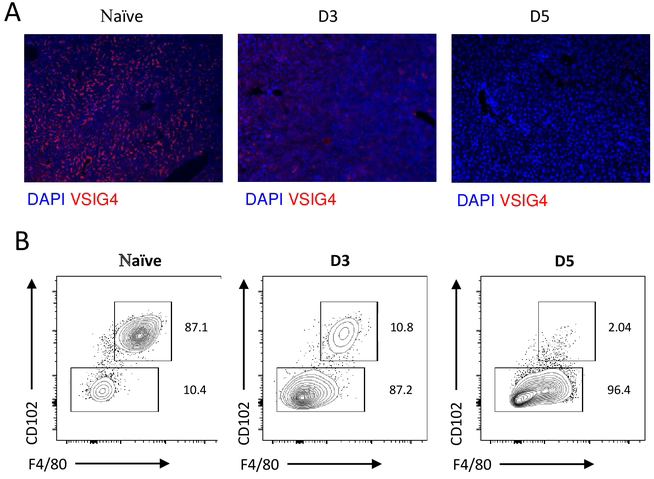
Systemic loss of TRMs during *T. gondii* infection. (A) Representative immunohistochemistry images showing the expression of VSIG4 in liver on days 3 and 5 post intraperitoneal infection with 20 cysts of ME49 *T. gondii*. (B) Flow cytometric analysis of large (CD11b+F4/80+CD102+) and small (CD11b+F4/80-CD102-) peritoneal macrophages measured on days 3 and 5 post intraperitoneal infection with *T. gondii.* The results are representative of three independent experiments.

**Supplemental Figure 3.**
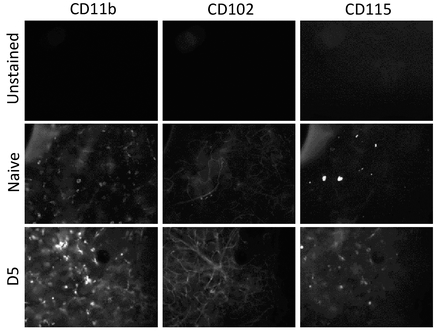
Analysis of LPMs migration to the omentum during *T. gondii* infection. WT (C57BL/6) mice were either uninfected (Naïve) or infected intraperitoneally with 20 cysts of the ME49 strain of *T. gondii* for 5 days (D5). Entire omenta from the infected and controlled mice were analyzed by whole mount staining for the presence of CD11b, CD102, and CD115 expressing cells.

## References

1 Gordon, S. & Taylor, P. R. Monocyte and macrophage heterogeneity. Nat Rev Immunol 5, 953–964, doi:10.1038/nri1733 (2005).

2 Pollard, J. W. Trophic macrophages in development and disease. Nat Rev Immunol 9, 259–270, doi:10.1038/nri2528 (2009).

3 Price, J. V. & Vance, R. E. The macrophage paradox. Immunity 41, 685–693, doi:10.1016/j.immuni.2014.10.015 (2014).

4 Austermann, J., Roth, J. & Barczyk-Kahlert, K. The Good and the Bad: Monocytes’ and Macrophages’ Diverse Functions in Inflammation. Cells 11, doi:10.3390/cells11121979 (2022).

5 Wynn, T. A., Chawla, A. & Pollard, J. W. Macrophage biology in development, homeostasis and disease. Nature 496, 445–455, doi:10.1038/nature12034 (2013).

6 Okabe, Y. & Medzhitov, R. Tissue biology perspective on macrophages. Nat Immunol 17, 9–17, doi:10.1038/ni.3320 (2016).

7 Mosser, D. M. & Edwards, J. P. Exploring the full spectrum of macrophage activation. Nat Rev Immunol 8, 958–969, doi:10.1038/nri2448 (2008).

8 Sinha, S. K. et al. Local M-CSF (Macrophage Colony-Stimulating Factor) Expression Regulates Macrophage Proliferation and Apoptosis in Atherosclerosis. Arterioscler Thromb Vasc Biol 41, 220–233, doi:10.1161/atvbaha.120.315255 (2021).

9 Hamilton, J. A. Colony-stimulating factors in inflammation and autoimmunity. Nat Rev Immunol 8, 533–544, doi:10.1038/nri2356 (2008).

10 Guilliams, M. et al. Dendritic cells, monocytes and macrophages: a unified nomenclature based on ontogeny. Nat Rev Immunol 14, 571–578, doi:10.1038/nri3712 (2014).

11 Blériot, C., Chakarov, S. & Ginhoux, F. Determinants of Resident Tissue Macrophage Identity and Function. Immunity 52, 957–970, doi:10.1016/j.immuni.2020.05.014 (2020).

12 Merad, M. et al. Langerhans cells renew in the skin throughout life under steady-state conditions. Nat Immunol 3, 1135–1141, doi:10.1038/ni852 (2002).

13 Yona, S. et al. Fate mapping reveals origins and dynamics of monocytes and tissue macrophages under homeostasis. Immunity 38, 79–91, doi:10.1016/j.immuni.2012.12.001 (2013).

14 Gomez Perdiguero, E., et al. Tissue-resident macrophages originate from yolk-sac-derived erythro-myeloid progenitors. Nature 518, 547–551, doi:10.1038/nature13989 (2015).

15 Hashimoto, D. et al. Tissue-resident macrophages self-maintain locally throughout adult life with minimal contribution from circulating monocytes. Immunity 38, 792–804, doi:10.1016/j.immuni.2013.04.004 (2013).

16 Schulz, C. et al. A lineage of myeloid cells independent of Myb and hematopoietic stem cells. Science 336, 86–90, doi:10.1126/science.1219179 (2012).

17 Bain, C. C. et al. Long-lived self-renewing bone marrow-derived macrophages displace embryo-derived cells to inhabit adult serous cavities. Nat Commun 7, ncomms11852, doi:10.1038/ncomms11852 (2016).

18 Hume, D. A. & MacDonald, K. P. Therapeutic applications of macrophage colony-stimulating factor-1 (CSF-1) and antagonists of CSF-1 receptor (CSF-1R) signaling. Blood 119, 1810–1820, doi:10.1182/blood-2011-09-379214 (2012).

19 Rojo, R. et al. Deletion of a Csf1r enhancer selectively impacts CSF1R expression and development of tissue macrophage populations. Nat Commun 10, 3215, doi:10.1038/s41467-019-11053-8 (2019).

20 MacDonald, K. P. et al. An antibody against the colony-stimulating factor 1 receptor depletes the resident subset of monocytes and tissue- and tumor-associated macrophages but does not inhibit inflammation. Blood 116, 3955–3963, doi:10.1182/blood-2010-02-266296 (2010).

21 Dai, X. M. et al. Targeted disruption of the mouse colony-stimulating factor 1 receptor gene results in osteopetrosis, mononuclear phagocyte deficiency, increased primitive progenitor cell frequencies, and reproductive defects. Blood 99, 111–120, doi:10.1182/blood.v99.1.111 (2002).

22 Solinas, G., Germano, G., Mantovani, A. & Allavena, P. Tumor-associated macrophages (TAM) as major players of the cancer-related inflammation. J Leukoc Biol 86, 1065–1073, doi:10.1189/jlb.0609385 (2009).

23 Jakubzick, C. V., Randolph, G. J. & Henson, P. M. Monocyte differentiation and antigen-presenting functions. Nat Rev Immunol 17, 349–362, doi:10.1038/nri.2017.28 (2017).

24 Martinez-Pomares, L. & Gordon, S. CD169+ macrophages at the crossroads of antigen presentation. Trends Immunol 33, 66–70, doi:10.1016/j.it.2011.11.001 (2012).

25 Park, J. & Hunter, C. A. The role of macrophages in protective and pathological responses to Toxoplasma gondii. Parasite Immunol 42, e12712, doi:10.1111/pim.12712 (2020).

26 Liddiard, K., Rosas, M., Davies, L. C., Jones, S. A. & Taylor, P. R. Macrophage heterogeneity and acute inflammation. Eur J Immunol 41, 2503–2508, doi:10.1002/eji.201141743 (2011).

27 Barth, M. W., Hendrzak, J. A., Melnicoff, M. J. & Morahan, P. S. Review of the macrophage disappearance reaction. J Leukoc Biol 57, 361–367, doi:10.1002/jlb.57.3.361 (1995).

28 Ghosn, E. E. et al. Two physically, functionally, and developmentally distinct peritoneal macrophage subsets. Proc Natl Acad Sci U S A 107, 2568–2573, doi:10.1073/pnas.0915000107 (2010).

29 Yarovinsky, F. Innate immunity to Toxoplasma gondii infection. Nat Rev Immunol 14, 109–121, doi:10.1038/nri3598 (2014).

30 Frickel, E. M. & Hunter, C. A. Lessons from Toxoplasma: Host responses that mediate parasite control and the microbial effectors that subvert them. J Exp Med 218, doi:10.1084/jem.20201314 (2021).

31 Araujo, A. et al. IFN-γ mediates Paneth cell death via suppression of mTOR. Elife 10, doi:10.7554/eLife.60478 (2021).

32 Leng, J., Butcher, B. A. & Denkers, E. Y. Dysregulation of macrophage signal transduction by Toxoplasma gondii: past progress and recent advances. Parasite Immunol 31, 717–728, doi:10.1111/j.1365-3024.2009.01122.x (2009).

33 Saeij, J. P. & Frickel, E. M. Exposing Toxoplasma gondii hiding inside the vacuole: a role for GBPs, autophagy and host cell death. Curr Opin Microbiol 40, 72–80, doi:10.1016/j.mib.2017.10.021 (2017).

34 Dunay, I. R., Fuchs, A. & Sibley, L. D. Inflammatory monocytes but not neutrophils are necessary to control infection with Toxoplasma gondii in mice. Infect Immun 78, 1564–1570, doi:10.1128/iai.00472-09 (2010).

35 Pollard, A. M., Knoll, L. J. & Mordue, D. G. The role of specific Toxoplasma gondii molecules in manipulation of innate immunity. Trends Parasitol 25, 491–494, doi:10.1016/j.pt.2009.07.009 (2009).

36 Goldszmid, R. S. et al. NK cell-derived interferon-γ orchestrates cellular dynamics and the differentiation of monocytes into dendritic cells at the site of infection. Immunity 36, 1047–1059, doi:10.1016/j.immuni.2012.03.026 (2012).

37 Askenase, M. H. et al. Bone-Marrow-Resident NK Cells Prime Monocytes for Regulatory Function during Infection. Immunity 42, 1130–1142, doi:10.1016/j.immuni.2015.05.011 (2015).

38 Grainger, J. R. et al. Inflammatory monocytes regulate pathologic responses to commensals during acute gastrointestinal infection. Nat Med 19, 713–721, doi:10.1038/nm.3189 (2013).

39 Neal, L. M. & Knoll, L. J. Toxoplasma gondii profilin promotes recruitment of Ly6Chi CCR2+ inflammatory monocytes that can confer resistance to bacterial infection. PLoS Pathog 10, e1004203, doi:10.1371/journal.ppat.1004203 (2014).

40 Robben, P. M., LaRegina, M., Kuziel, W. A. & Sibley, L. D. Recruitment of Gr-1+ monocytes is essential for control of acute toxoplasmosis. J Exp Med 201, 1761–1769, doi:10.1084/jem.20050054 (2005).

41 Del Rio, L. et al. Toxoplasma gondii triggers myeloid differentiation factor 88-dependent IL-12 and chemokine ligand 2 (monocyte chemoattractant protein 1) responses using distinct parasite molecules and host receptors. J Immunol 172, 6954–6960, doi:10.4049/jimmunol.172.11.6954 (2004).

42 Safronova, A. et al. Alarmin S100A11 initiates a chemokine response to the human pathogen Toxoplasma gondii. Nat Immunol 20, 64–72, doi:10.1038/s41590-018-0250-8 (2019).

43 Sturge, C. R. & Yarovinsky, F. Complex immune cell interplay in the gamma interferon response during Toxoplasma gondii infection. Infect Immun 82, 3090–3097, doi:10.1128/iai.01722-14 (2014).

44 Yamamoto, M. et al. A cluster of interferon-γ-inducible p65 GTPases plays a critical role in host defense against Toxoplasma gondii. Immunity 37, 302–313, doi:10.1016/j.immuni.2012.06.009 (2012).

45 Collazo, C. M. et al. Inactivation of LRG-47 and IRG-47 reveals a family of interferon gamma-inducible genes with essential, pathogen-specific roles in resistance to infection. J Exp Med 194, 181–188, doi:10.1084/jem.194.2.181 (2001).

46 Denkers, E. Y. From cells to signaling cascades: manipulation of innate immunity by Toxoplasma gondii. FEMS Immunol Med Microbiol 39, 193–203, doi:10.1016/s0928-8244(03)00279-7 (2003).

47 Murray, H. W., Spitalny, G. L. & Nathan, C. F. Activation of mouse peritoneal macrophages in vitro and in vivo by interferon-gamma. J Immunol 134, 1619–1622 (1985).

48 Taylor, G. A., Feng, C. G. & Sher, A. p47 GTPases: regulators of immunity to intracellular pathogens. Nat Rev Immunol 4, 100–109, doi:10.1038/nri1270 (2004).

49 Raetz, M. et al. Parasite-induced TH1 cells and intestinal dysbiosis cooperate in IFN-γ-dependent elimination of Paneth cells. Nat Immunol 14, 136–142, doi:10.1038/ni.2508 (2013).

50 Cohen, S. B. & Denkers, E. Y. The gut mucosal immune response to Toxoplasma gondii. Parasite Immunol 37, 108–117, doi:10.1111/pim.12164 (2015).

51 Langer, V. et al. IFN-γ drives inflammatory bowel disease pathogenesis through VE-cadherin-directed vascular barrier disruption. J Clin Invest 129, 4691–4707, doi:10.1172/jci124884 (2019).

52 Platanias, L. C. Mechanisms of type-I- and type-II-interferon-mediated signalling. Nat Rev Immunol 5, 375–386, doi:10.1038/nri1604 (2005).

53 Hu, X. & Ivashkiv, L. B. Cross-regulation of signaling pathways by interferon-gamma: implications for immune responses and autoimmune diseases. Immunity 31, 539–550, doi:10.1016/j.immuni.2009.09.002 (2009).

54 Rivera-Cuevas, Y. et al. Toxoplasma gondii exploits the host ESCRT machinery for parasite uptake of host cytosolic proteins. PLoS Pathog 17, e1010138, doi:10.1371/journal.ppat.1010138 (2021).

55 Wang, Y. et al. Genome-wide screens identify Toxoplasma gondii determinants of parasite fitness in IFNγ-activated murine macrophages. Nat Commun 11, 5258, doi:10.1038/s41467-020-18991-8 (2020).

56 Wang, Y. et al. Three Toxoplasma gondii Dense Granule Proteins Are Required for Induction of Lewis Rat Macrophage Pyroptosis. mBio 10, doi:10.1128/mBio.02388-18 (2019).

57 Huang, Z. et al. The intrinsically disordered protein TgIST from Toxoplasma gondii inhibits STAT1 signaling by blocking cofactor recruitment. Nat Commun 13, 4047, doi:10.1038/s41467-022-31720-7 (2022).

58 Panas, M. W. & Boothroyd, J. C. Seizing control: How dense granule effector proteins enable Toxoplasma to take charge. Mol Microbiol 115, 466–477, doi:10.1111/mmi.14679 (2021).

59 Barrat, F. J., Crow, M. K. & Ivashkiv, L. B. Interferon target-gene expression and epigenomic signatures in health and disease. Nat Immunol 20, 1574–1583, doi:10.1038/s41590-019-0466-2 (2019).

60 Christian, D. A., et al. cDC1 Coordinate Innate and Adaptive Responses in the Omentum required for T cell Priming and Memory. bioRxiv, 2020.2007.2021.214809, doi:10.1101/2020.07.21.214809 (2020).

61 Gautier, E. L. et al. Gene-expression profiles and transcriptional regulatory pathways that underlie the identity and diversity of mouse tissue macrophages. Nat Immunol 13, 1118–1128, doi:10.1038/ni.2419 (2012).

62 Kim, K. W. et al. MHC II+ resident peritoneal and pleural macrophages rely on IRF4 for development from circulating monocytes. J Exp Med 213, 1951–1959, doi:10.1084/jem.20160486 (2016).

63 McCann, K. J. et al. IFNγ regulates NAD+ metabolism to promote the respiratory burst in human monocytes. Blood Adv 6, 3821–3834, doi:10.1182/bloodadvances.2021005776 (2022).

64 Sanin, D. E. et al. A common framework of monocyte-derived macrophage activation. Sci Immunol 7, eabl7482, doi:10.1126/sciimmunol.abl7482 (2022).

65 Weichhart, T., Hengstschläger, M. & Linke, M. Regulation of innate immune cell function by mTOR. Nat Rev Immunol 15, 599–614, doi:10.1038/nri3901 (2015).

66 Su, X. et al. Interferon-γ regulates cellular metabolism and mRNA translation to potentiate macrophage activation. Nat Immunol 16, 838–849, doi:10.1038/ni.3205 (2015).

67 Guilliams, M. et al. Spatial proteogenomics reveals distinct and evolutionarily conserved hepatic macrophage niches. Cell 185, 379–396.e338, doi:10.1016/j.cell.2021.12.018 (2022).

68 Zhou, X. et al. Microenvironmental sensing by fibroblasts controls macrophage population size. Proc Natl Acad Sci U S A 119, e2205360119, doi:10.1073/pnas.2205360119 (2022).

69 Smith, A. M. et al. M-CSF increases proliferation and phagocytosis while modulating receptor and transcription factor expression in adult human microglia. J Neuroinflammation 10, 85, doi:10.1186/1742-2094-10-85 (2013).

70 Han, S. et al. Macrophage-colony stimulating factor enhances MHC-restricted presentation of exogenous antigen in dendritic cells. Cytokine 32, 187–193, doi:10.1016/j.cyto.2005.08.002 (2005).

71 Curry, J. M. et al. M-CSF signals through the MAPK/ERK pathway via Sp1 to induce VEGF production and induces angiogenesis in vivo. PLoS One 3, e3405, doi:10.1371/journal.pone.0003405 (2008).

72 Sehgal, A., Irvine, K. M. & Hume, D. A. Functions of macrophage colony-stimulating factor (CSF1) in development, homeostasis, and tissue repair. Semin Immunol 54, 101509, doi:10.1016/j.smim.2021.101509 (2021).

73 Ushach, I. & Zlotnik, A. Biological role of granulocyte macrophage colony-stimulating factor (GM-CSF) and macrophage colony-stimulating factor (M-CSF) on cells of the myeloid lineage. J Leukoc Biol 100, 481–489, doi:10.1189/jlb.3RU0316-144R (2016).

74 Becker, A. M., Walcheck, B. & Bhattacharya, D. ADAM17 limits the expression of CSF1R on murine hematopoietic progenitors. Exp Hematol 43, 44–52.e41-43, doi:10.1016/j.exphem.2014.10.001 (2015).

75 Vahidi, A., Glenn, G. & van der Geer, P. Identification and mutagenesis of the TACE and γ-secretase cleavage sites in the colony-stimulating factor 1 receptor. Biochem Biophys Res Commun 450, 782–787, doi:10.1016/j.bbrc.2014.06.061 (2014).

76 Tang, J. et al. Neutrophil and Macrophage Cell Surface Colony-Stimulating Factor 1 Shed by ADAM17 Drives Mouse Macrophage Proliferation in Acute and Chronic Inflammation. Mol Cell Biol 38, doi:10.1128/mcb.00103-18 (2018).

77 Parks, W. C., Wilson, C. L. & López-Boado, Y. S. Matrix metalloproteinases as modulators of inflammation and innate immunity. Nat Rev Immunol 4, 617–629, doi:10.1038/nri1418 (2004).

78 Tcyganov, E. N. et al. Distinct mechanisms govern populations of myeloid-derived suppressor cells in chronic viral infection and cancer. J Clin Invest 131, doi:10.1172/jci145971 (2021).

